## Supplementary figures and images for "The *Aspergillus nidulans* velvet domain containing transcription factor VeA is shuttled from cytoplasm into nucleus during vegetative growth and stays there for sexual development, but has to return into cytoplasm for asexual development"

### Supplementary figures

Fig S1_Strohdiek *et al*.

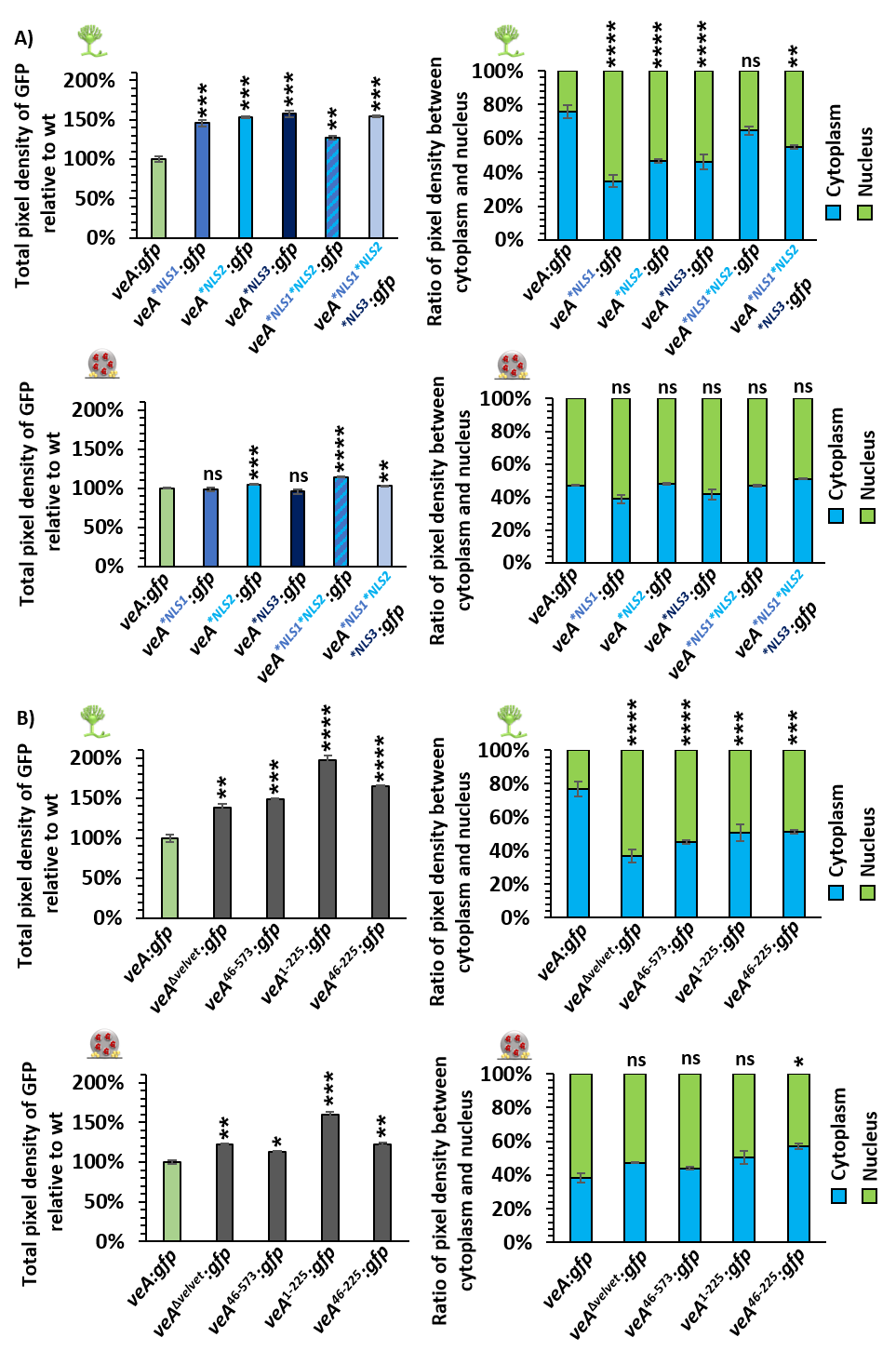

Fig S2_Strohdiek *et al*.

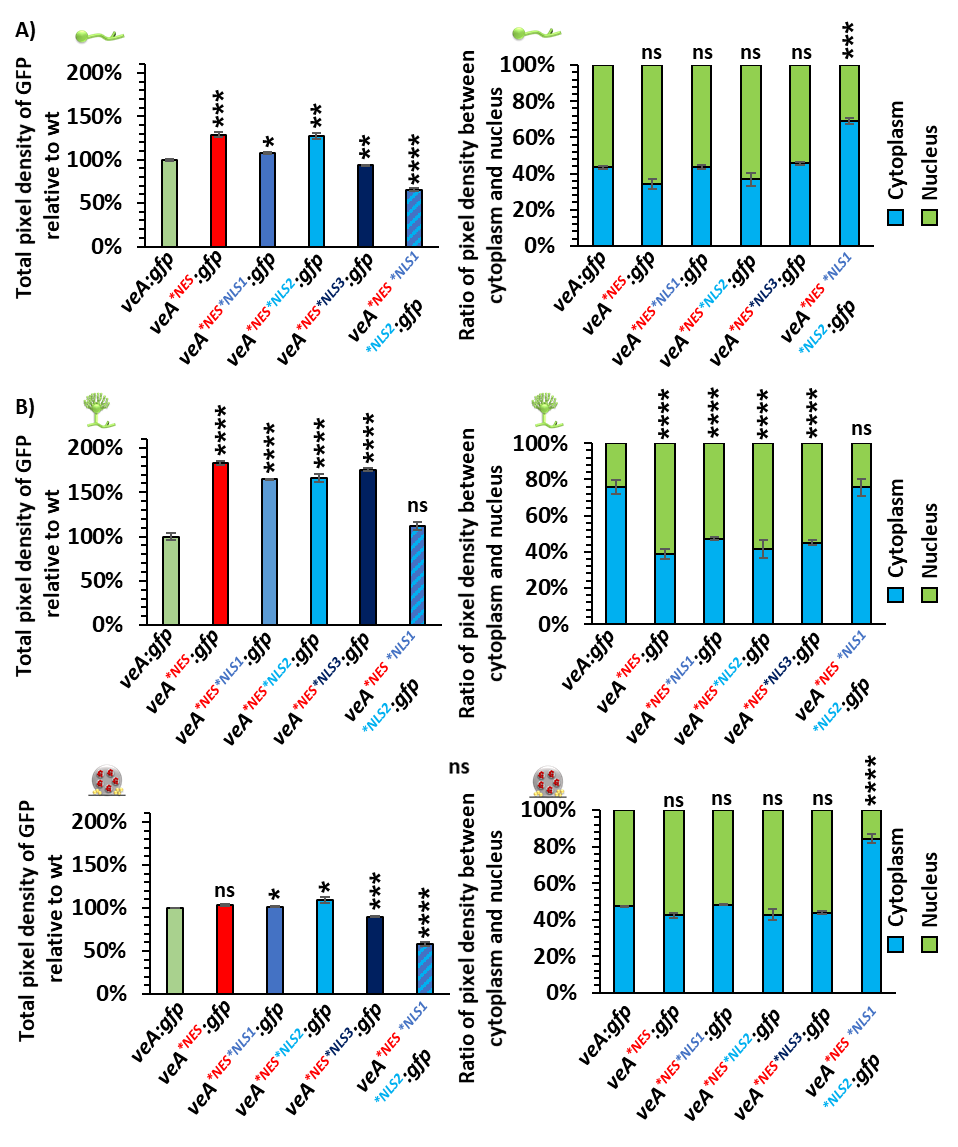

Fig S3_Strohdiek *et al*.

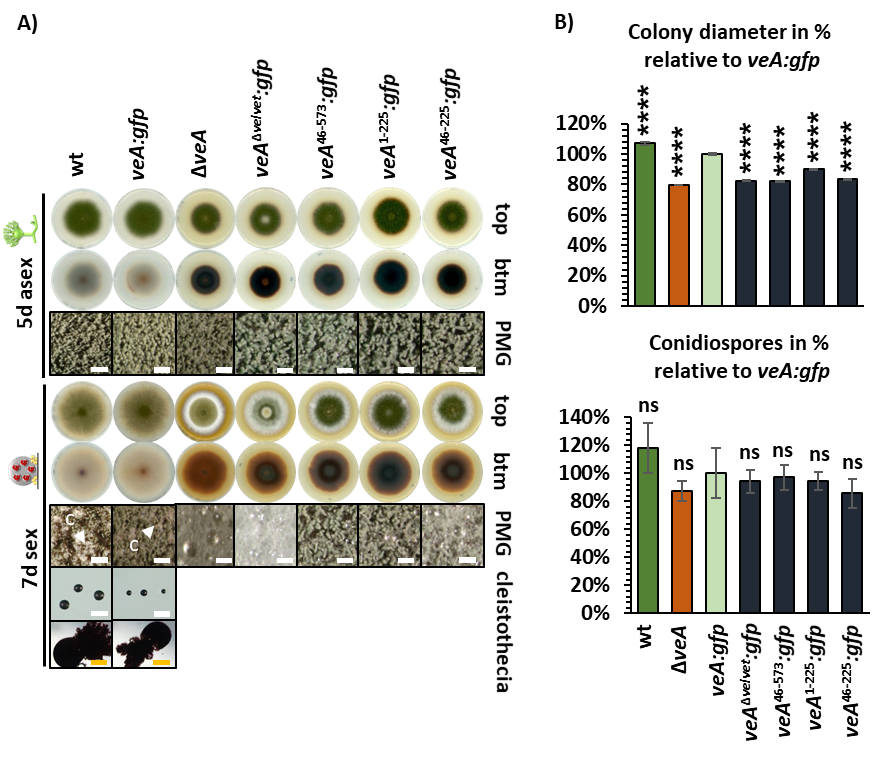

Fig S4_Strohdiek *et al*.

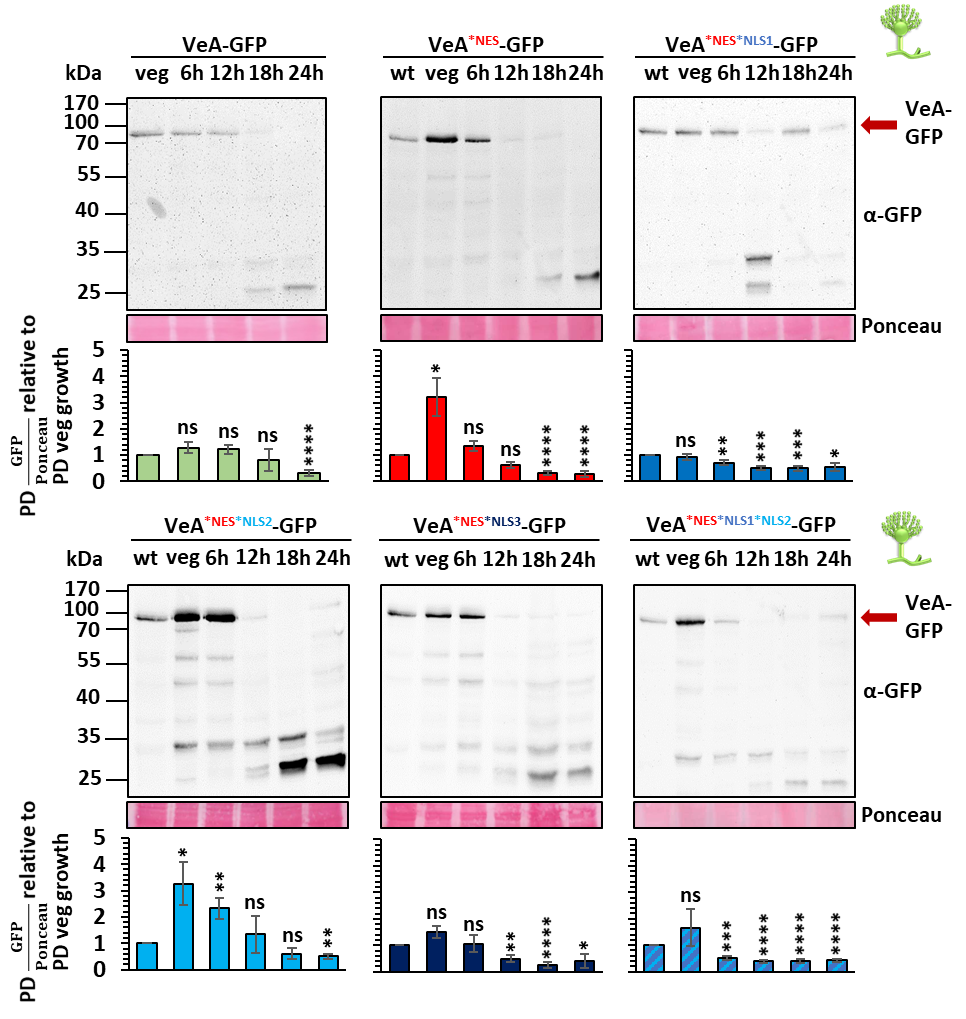
