## Supplementary tables for "The *Aspergillus nidulans* velvet domain containing transcription factor VeA is shuttled from cytoplasm into nucleus during vegetative growth and stays there for sexual development, but has to return into cytoplasm for asexual development"

Table S1

| **No.** | **Chemical name** | **Molecular Formula** | **Calc. exact mass [M]** | **Measured exact mass [M+H]^+^** | **Rt (min)** | **Confirmed by** | **Reference** | **Gene cluster** |
| --- | --- | --- | --- | --- | --- | --- | --- | --- |
| I | Cichorine | C_10_H_11_NO_3_ | 193.073894 | 194.0814 | 7.6 | A, C | Sanchez et al., 2012 | cic, CicF (nrPKS, AN6448) |
| II | F-9775A/B | C_21_H_16_O_8_ | 396.08452 | 397.0917 | 9.5,  10.5 | A, C | Bok et al., 2009 | ors, OrsA (nrPKS, AN7909) |
| III | Austinol | C_25_H_40_O_8_ | 458.19407 | 459.2012 | 12.7 | A, B | Lo et al., 2012 | aus, AusA (nrPKS, AN8383) |
| IV | Dehydroaustinol | C_25_H_28_O_8_ | 456.17842 | 457.1853 | 13.1 | A, B | Lo et al., 2012 | aus, AusA (nrPKS, AN8383) |
| V | Sterigmatocystin | C_18_H_12_O_6_ | 324.06339 | 325.0703 | 15.3 | A,B | Yu and Leonard, 1995 | stc, StcA (nrPKS, AN7825) |
| VI | Emericellin | C_25_H_28_O_5_ | 408.193675 | 409.2005 391.1898  [M-H_2_O+H]^+^ | 24.6 | A, B | Sanchez et al., 2011 | mdp, mdpG (nrPKS, AN0150) |
| VII | Shamixanthone | C_25_H_26_O_5_ | 406.178025 | 389.1743  [M-H_2_O+H]^+^ | 24.8 | A, B | Sanchez et al., 2011 | mdp, MdpG (nrPKS, AN0150) |
| VIII | Epishamixanthone | C_25_H_26_O_5_ | 406.178025 | 389.1741  [M-H_2_O+H]^+^ | 25.8 | A, B | Sanchez et al., 2011 | mdp, MdpG (nrPKS, AN0150) |

Peak numbers from CAD and EIC (Fig 4, Fig 5) correspond to secondary metabolites. A: Exact mass measurement, B: retention time (8,37)/ comparison with commercial standards for sterigmatocystin, C: UV/VIS spectrum (55,56).

Table S2

| Plasmid | Genotype | Reference |
| --- | --- | --- |
| pME4319 | Self-excising β-rec/six phleo-RM containing vector | Dr. J. Gerke, p.c. |
| pME5533 | *veA^1-225^*, C-terminal *gfp*, phleo cassette | This study |
| pME5534 | *veA^46-225^*, C-terminal *gfp*, phleo cassette | This study |
| pME5544 | *veA^L188A/L192A^,* C-terminal *gfp,* phleo cassette | This study |
| pME5546 | *veA^L188A/L192A/K509A/K510A^,* C-terminal *gfp*, phleo cassette | This study |
| pME5547 | *veA^L188A/L192A/R540A/R541A/K559A/R560A^,* C-terminal *gfp,* phleo cassette | This study |
| pME5548 | *veA^K28A/K29A/K41A/R42A/L188A/L192A^,* C-terminal *gfp,* phleo cassette | This study |
| pME5549 | *veA^K28A/K29A/K41A/R42A/L188A/L192AK509A/K510A^*, C-terminal *gfp*, phleo cassette | This study |
| pME5550 | *veA^K28A/K29A/K41A/R42A/L188A/L192A/R540A/R541A/K559A/R560A^,* C-terminal *gfp*, phleo cassette | This study |
| pME5551 | *veA^K28A/K29A/K41A/R42A//L188A/L192A/K509A/K510A/R540A/R541A/K559A/R560A^,* C-terminal *gfp*, phleo cassette | This study |
| pME5553 | *veA^K28A/K29A/K41A/R42A/K509A/K510A^*, C-terminal *gfp*, phleo cassette | This study |
| pME5554 | *veA^K509A/K510A^,* C-terminal *gfp*, phleo cassette | This study |
| pME5555 | *veA^R540A/R541A/K559A/R560A^,* C-terminal *gfp*, phleo cassette | This study |
| pME5557 | *veA^R540A/R541A/K559A/R560A/R540A/R541A/K559A/R560A^,* C-terminal *gfp,* phleo cassette | This study |

Table S3

| Strain | Genotype | Reference |
| --- | --- | --- |
| AGB552 | ∆*nkuA::argB, pabaA1, yA2, veA^+^* | Bayram *et al*., 2012 |
| AGB1195 | ∆*nkuA::argB*, *pyroA4, pyrG89*, *veA^+^*, ∆*vosA*, *velB:gfp:six*, ∆*veA* | Dr. S. Thieme, p.c. |
| AGB1337 | ∆*nkuA::argB*, ∆*veA::six*, *pabaA1, rfp:h2A, nat^R^* | Dr. J. Gerke, p.c |
| AGB1346 | *∆nkuA, ^P^veA:veA^K28A/K29A/K41A/R42A^:gfp:six:veA^T^; pabaA1; rfp:h2A, phleo^R^* | Dr. J. Gerke, p.c |
| AGB1351 | *∆nkuA::argB, ^P^veA:veA:gfp:six:veA^T^; pabaA1; rfp:h2A, phleo^R^* | Dr. J. Gerke, p.c |
| AGB1353 | *∆nkuA::argB, pabaA1, ^P^veA:veA^46-537^:gfp:six:veA^T^, rfp:h2A, phleo^R^* | Dr. J. Gerke p.c. |
| AGB1358 | *∆nkuA::argB, pabaA1, rfp:h2A, nat^R^* | Dr. J. Gerke, p.c |
| AGB1392 | *∆nkuA::argB, pabaA1, veA^∆velvet^:gfp, rfp:h2A, nat^R^* | Dr. J. Gerke, p.c. |
| AGB1666 | ∆*nkuA::argB, ^P^veA:veA^1-225^:L:gfp:six:veA^T^* | This study |
| AGB1667 | ∆*nkuA::argB, ^P^veA:veA^46-225^:L:gfp:six:veA^T^* | This study |
| AGB1677 | *∆nkuA::argB, ^P^veA:veA^L188A/L192A^:gfp:six:veA^T^; pabaA1; rfp:h2A* | This study |
| AGB1678 | *∆nkuA::argB, ^P^veA:veA^L188A/L192A/K509A/K510A^:gfp:six:veA^T^; pabaA1; rfp:h2A* | This study |
| AGB1679 | *∆nkuA::argB, ^P^veA:veA^L188A/L192A/R540A/R541A/K559A/R560A^:gfp:six:veA^T^; pabaA1; rfp:h2A* | This study |
| AGB1680 | *∆nkuA, ^P^veA:veA^K28A/K29A/K41A/R42A/L188A/L192A^:gfp:six:veA^T^; pabaA1; rfp:h2A* | This study |
| AGB1681 | *∆nkuA::argB, ^P^veA:veA^K28A/K29A/K41A/R42A/L188A/L192AK509A/K510A^:gfp:six:veA^T^; pabaA1; rfp:h2A* | This study |
| AGB1690 | *∆nkuA::argB, ^P^veA:veA^K28A/K29A/K41A/R42A/K509A/K510A^:gfp:six:veA^T^; pabaA1; rfp:h2A* | This study |
| AGB1691 | *∆nkuA::argB, ^P^veA:veA^K509A/K510A^:gfp:six:veA^T^; pabaA1; rfp:h2A* | This study |
| AGB1692 | *∆nkuA::argB, ^P^veA:veA^R540A/R541A/K559A/R560A^:gfp:six:veA^T^; pabaA1; rfp:h2A* | This study |
| AGB1695 | *∆nkuA::argB, ^P^veA:veA^K28A/K29A/K41A/R42A^:gfp:six:veA^T^; pabaA1; rfp:h2A* | This study |
| AGB1700 | *∆nkuA::argB, ^P^veA:veA^K28A/K29A/K41A/R42A/K509A/K510A/R540A/R541A/K559A/R560A^:gfp:six:veA^T^; pabaA1; rfp:h2A* | This study |

^P^: promoter, ^T^: terminator, ^R^: resistance, PP: PreScission cleavage site; L: linker, nat^R^: nat resistance, phleo^R^: phleo resistance.

Table S4

| Primer | Sequence | Reference |
| --- | --- | --- |
| oAS35 | aggaattcgatATTTGTTTAAACTTAACCTAAATGCACGCACT | This study |
| oAS36 | CTTGATGGGATAACACAAAATGCT | This study |
| oAS37 | ACCACCGCTACCACCGCAGCCCTGCTCCGCAA | This study |
| oAS38 | GGTGGTAGCGGTGGTGTG | This study |
| oAS39 | ataggcctgagATTTCTACTTGTACAGTTCGTCCATG | This study |
| oAS40 | atatggccatctCACAAGAATTCTGCCGGCGTTTATTT | This study |
| oAS41 | gataagcttgatCACGTTTAAACGGATTGAATGCGGACG | This study |
| oAS42 | TGTTATCCCATCAAGatgTGTGGTCAAGGTTCGAAAT | This study |
| oAS111 | CATCGAGAAGAGAAGATCTTTCATAGGACGGGGCGGCCTGCAT | This study |
| oAS112 | CTTCTCTTCTCGATGGCCCCGACCAGATGGCTTATAAGGCCGCCAACGGGC | This study |
| oAS127 | GTTCGAGATGACCTCCGCCCGGAACTCCCGCAATTCTTGCGGTGAC | This study |
| oAS128 | GTCACCGCAAGAATTGCGGGAGTTCCGGGCGGAGGTCATCTCGAAC | This study |
| oAS129 | CGACTCTTCGTACGTCGCCGCCGACAACATCGAGTT | This study |
| oAS130 | AACTCGATGTTGTCGGCGGCGACGTACGAAGAGTCG | This study |
| oAS135 | ACCACAAGCTCTCGCGgcCgcGGGCTGCTGCATAAT | This study |
| oAS136 | ATTATGCAGCAGCCCgcGgcCGCGAGAGCTTGTGGT | This study |
| oAS137 | ATCACTCGCGAGGGAgcGgcGATTACCTATAAATTG | This study |
| oAS138 | CAATTTATAGGTAATCgcCgcTCCCTCGCGAGTGAT | This study |
| oAS139 | GTTCGAGATGACCTCCAGCCGGAACTCCAAGAATTCTTGCGGTGAC | This study |
| oAS140 | GTCACCGCAAGAATTCTTGGAGTTCCGGCTGGAGGTCATCTCGAAC | This study |

Table S5

| Primer | Sequence | Reference |
| --- | --- | --- |
| JG814 | CAACGAGCATCAGCACAAACAT | Dr. J. Gerke, p.c |
| JG815 | ATCTTCTACGCCGATTCCTGCT | Dr. J. Gerke, p.c |
| JG880 | ACGACGAGGGTCTTATTGTCA | Dr. J. Gerke, p.c |
| JG881 | CCTTCTCCTGGGTAGTGAA | Dr. J. Gerke, p.c |
| MG277 | CGTCTTCTTCGCAAGGGAAACT | Dr. M. Gulko, p.c |
| MG278 | CGGGTTTTCTTGTTGTCACGAG | Dr. M. Gulko, p.c |

Table S6

|  | Velvet domain of VeA |
| --- | --- |
| PDB id | 9I2S |
| Wavelength [Å] | 0.8 |
| Resolution range [Å] | 47.21    - 2.4 (2.486    - 2.4) |
| Space group | P 2(1) 2(1) 2 |
| Unit cell [Å, °] | 106.04 156.08 52.73 90 90 90 |
| Total reflections | 144071 (15106) |
| Unique reflections | 34475 (3372) |
| Multiplicity | 4.2 (4.4) |
| Completeness (%) | 98.00 (98.2) |
| Mean I/sigma(I) | 22.93 (1.05) |
| Wilson B-factor [Å^2^] | 71.49 |
| R-merge | 0.041 (1.821) |
| R-meas | 0.0470 (2.078) |
| R-pim | 0.02504 (1.008) |
| CC1/2 | 1 (0.504) |
| CC* | 1 (0.819) |
| Reflections used in refinement | 34383 (3370) |
| Reflections used for R-free | 1719 (169) |
| R-work | 0.2193 (0.4096) |
| R-free | 0.2531 (0.4813) |
| CC(work) | 0.957 (0.499) |
| CC(free) | 0.963 (0.316) |
| Number of non-hydrogen atoms | 5530 |
| macromolecules | 5486 |
| ligands | 0 |
| solvent | 44 |
| Protein residues | 682 |
| RMS (bonds) [Å] | 0.004 |
| RMS (angles) [°] | 0.70 |
| Ramachandran favored (%) | 97.87 |
| Ramachandran allowed (%) | 2.13 |
| Ramachandran outliers (%) | 0.00 |
| Rotamer outliers (%) | 1.80 |
| Clashscore | 5.78 |
| Average B-factor [Å^2^] | 102.05 |
| macromolecules | 102.28 |
| solvent | 74.53 |
| Number of TLS groups | 26 |

Statistics for the highest-resolution shell are shown in parentheses.
